# Molecular Insights into Long COVID: Plasma Proteomics Reveals Oxidative Stress, Coagulation Cascade Activation, and Glycolytic Imbalance

**DOI:** 10.1101/2025.07.16.665198

**Authors:** Mohammad Mobarak H Chowdhury, Akouavi Julite Irmine Quenum, Christine Rioux-Perreault, Jean-François Lucier, Subburaj Ilangumaran, Alain Piché, Hugues Allard-Chamard, Sheela Ramanathan

## Abstract

Persistent symptoms following SARS-CoV-2 infection are the hallmark of post-COVID condition (PCC), also referred to as long COVID. However, our knowledge is limited on the underlying molecular mechanisms. In this study, we performed data-independent acquisition mass spectrometry plasma proteomics (DIA-MS) to identify molecular alterations associated with PCC. DIA-MS proteomic analysis revealed a few proteins linked to oxidative stress that had altered expression. Notably, PCC samples exhibited downregulation of the antioxidant protein peroxiredoxin 6 (PRDX6) and upregulation of oxidative stress-associated proteins particularly vanin-1 (VNN1) and paraoxonase-3 (PON3). Additionally, the PCC group showed significantly higher levels of six proteins (PCSK9, CST3, C1Q, CPB2, KNG1 GAPDH), which were linked to pathways involving glycolysis, complement and coagulation cascades, and inflammation. Oxidative stress analysis confirmed that PCC samples had significantly higher levels of DNA damage (8-OHdG) than the convalescent group, whereas antioxidant markers, such as reduced and oxidized glutathione (GSH and GSSG), were significantly lower in PCC samples than in uninfected controls. Our observations point towards ongoing oxidative and inflammatory processes in PCC and suggest potential targets for biomarker development and therapeutic intervention.

## Introduction

Even though the majority of COVID-19 infections are relatively mild with recovery typically occurring within 2–3 weeks [1, 2], a significant proportion of COVID-19 patients experience a panoply of long-lasting symptoms after the acute SARS-CoV-2 infection. These symptoms, collectively known as “post COVID-19 condition” (**PCC**, or “long COVID”), compromise their health-related quality of life and impair their daily functioning [3–6]. PCC is estimated to affect 65 million individuals worldwide [7, 8], including children [9–12], lasts for months or even years [13]. PCC encompasses a spectrum of clinical symptoms affecting multiple organs that persist for more than 3 months after the resolution of the initial infection [9, 14–20]. They include, but are not restricted to, dysautonomia affecting cardiac (palpitations, syncope, dysrhythmias and postural symptoms), neuropsychiatric (insomnia, chronic headache, brain fog, defects in memory, mood impairment and pain syndromes), and respiratory (dyspnea and cough) [7, 14, 21], as well as asthenia.

PCC has notable impacts on public health, while the understanding of the exact molecular pathways behind its progression are yet poorly characterized. Nonetheless, oxidative stress and abnormal immune response are now recognized as important variables among several possible pathogenic components that may be crucial in controlling the chronic and persistent challenges observed in PCC individuals [22, 23]. Various studies have tried to identify biomarkers that could predict the development of PCC [24]. Lipid peroxidation, degradation of proteins, and DNA strand damage are all carried out by reactive oxygen species (ROS), which seriously impair cellular metabolism. It has been reported that Covid-19 non-survivors had a reduced level of key antioxidants, including vitamin E and glutathione (GSH), while survivors had unexpectedly lower superoxide dismutase (SOD) levels and elevated oxidation-associated protein product (AOPP), despite no significant variations in 8-OHdG and malondialdehyde (MDA) levels [25]. Furthermore, several studies have shown aggravated indicators of oxidative stress during acute COVID-19, in addition to raised levels of malondialdehyde (MDA), a lipid peroxidation marker, in individuals experiencing with post-COVID conditions (PCC) [22, 23]. Increased redox stress has been closely linked to several clinical problems, such as metabolic syndrome, pulmonary dysfunction, cardiovascular disease, and neurological disorders which are commonly observed in both severe COVID-19 and PCC [26]. It remains unclear whether the observed increase in reactive oxygen species (ROS) drives the clinical manifestations, or if infection or the underlying conditions, such as obesity, are themselves responsible for elevating ROS levels. In this study, we analyzed the plasma proteome of convalescent, PCC and uninfected individuals at 3 months after diagnosis of SARS-CoV-2 by PCR that had developed mild infection without hospitalization. Our results show that the changes in the plasma proteome persist even at three months after mild SARS-CoV-2 infection and suggest that persistence of oxidative stress may persist even after 3 months in recovered SARS-CoV-2 infected individuals.

## 2. Materials and Methods

### 2.1. Participants recruitment

This study used data and plasma samples from the Biobanque Québécoise de la COVID-19 (BQC19), an ongoing provincial healthcare initiative in Quebec, Canada, aimed at collecting biologic specimens, including blood cells and plasma, along with detailed anthropometric and clinical data from individuals diagnosed with SARS-CoV-2 infection confirmed through PCR testing. Since March 26, 2020, this initiative has been recruiting adult participants (aged 18 years and older) with varying degrees of disease severity from several healthcare centers throughout Quebec [27]. For this study, samples were obtained from one of the participating center – Centre de Recherche du CHUS – between March 2020 and October 2021, with all infections confirmed before October 2021. The study design, data collection and procedures have been described in detail elsewhere [28]. SARS-CoV-2 infection severity was classified based on the World Health Organization (WHO) criteria as asymptomatic, mild, moderate, or severe [29]. The study was conducted with approval from the ethics review board of the Centre de Recherche du Centre Hospitalier Universitaire de Sherbrooke (protocol # 2022-4415).

### 2.2 Application of Mass Spectrometry for Protein Analysis

#### 2.2.1. Methods for Protein Preparation and Protease Digestion Prior to Mass Spectrometry

Plasma samples were prepared for proteomic analysis by the proteomics platform at Université de Sherbrooke following established protocols as described in Chowdhury et al. [30].

#### 2.2.2. Analysis and visualization of PCA and differentially expressed proteomic data

Raw data files were processed using the DIA-NN software v.1.8.1 (DIA-NN: neural networks and interference correction enable deep proteome coverage in high throughput) [31] available on GitHub (https://github.com/vdemichev/DiaNN). DIA-NN was installed in an Apptainer container using the Docker image from Docker Hub (https://hub.docker.com/layers/biocontainers/diann/v1.8.1_cv1/images).The analysis was conducted with default parameters, allowing for up to two permissive cleavages and protein N-terminal methionine as a variable in the in-silico digestion.

The human proteome (FASTA file UP000005640) was obtained from the Uniprot database (https://ftp.uniprot.org/pub/databases/uniprot/current_release/knowledgebase/reference_proteomes/Euk aryota/UP000005640/), comprising 96,418 protein entries. For the FASTA-based searches, DIA-NN was configured to conduct in-silico digestion of the protein sequences. A mass tolerance of 20 ppm was applied for both MS1 precursor ions and MS2 fragment ions. The peptide length was limited to 7-30 amino acids, and precursor charge states were set between 1 and 5. The precursor m/z range was configured from 100 to 1700, and fragment m/z from 100 to 1500. These parameters were employed for both library generation through in-silico approaches and library-free searches. Match Between Runs (MBR) and smart profiling functionalities were activated to construct a spectral library directly from the DIA data. Carboxyamidomethylation (Unimod4) and methionine oxidation (Unimod35) were designated as fixed modifications, while N-terminal protein acetylation was treated as a variable modification. Post-analysis was conducted using R (version 4.3.2; www.R-project.org), resulting in the final output file titled “unique_genes_matrix.tsv.” This matrix contained quantified gene-specific peptides filtered using a global false discovery rate (FDR) of 1%, applying both global q-values for protein groups and global as well as run-specific q-values for precursors. Twenty repeated samples were utilized solely for validation purposes and were excluded from statistical analyses; consequently, data from 150 unique samples were included in the final analysis. These curated datasets, along with group comparisons, are available in the Supplementary Table. GraphPad Prism version 10.0.3 (GraphPad, Boston, MA, USA) was employed to create volcano plots and quantify protein abundance between the study groups. Venn diagrams were generated using the jvenn online tool (https://jvenn.toulouse.inrae.fr/app/index.html) and DIA-NN software. Furthermore, PCA was conducted to cluster peptide-based intensity and intact protein detection across three groups using ELISA. The analysis was performed with the SRplot web server (https://www.bioinformatics.com.cn/en).

#### 2.2.3. Protein Validation by ELISA Assays

Protein levels for GAPDH (Catalog # EH207RB), Proprotein Convertase 9/PCSK9 (Catalog # EH384RB), sCD26 (Catalog # BMS235), Carboxypeptidase B2/CPB2 (Catalog # EH68RB), and C1q (Catalog # BMS2099 or BMS2099TEN) in plasma were quantified according to the specific protocols provided by the manufacturers (ThermoFisher, USA). CST3 (Cystatin C) ELISA kit was obtained from Sinobiologicals (# sek10439).

#### 2.2.4. Gene ontology and pathway enrichment analyses

To explore the functional characteristics of common proteins between PCC, convalescent and uninfected groups, we performed the comprehensive enrichment analyses employing the SRplot web server, a robust web-based gene set enrichment tool (http://www.bioinformatics.com.cn/srplot). Visualization and further analysis of the resulting protein-protein interaction (PPI) networks were performed using Cytoscape software (version 3.8.2). Furthermore, we employed the cytoHubba plugin within Cytoscape for applying the degree of topological algorithm. This approach allowed us to select the most critical hub proteins with the highest degree of connectivity which are potentially significant in the underlying biological processes.

### 2.3. Antioxidant Glutathione (GSH+GSSG) measurement in Plasma by colorimetric Assay

Plasma samples isolated from fresh blood through centrifugation and stored at -80. The total levels of glutathione (GSH + GSSG) in plasma were quantified using the Glutathione Colorimetric Detection Kit (Invitrogen, Catalog Number: EIAGSHC). Samples were initially diluted with the Assay Buffer provided in the kit to ensure their concentrations were within the detectable range. To specifically measure GSSG levels, a subset of samples was treated with 2-vinylpyridine (2VP) in separate tubes to block free GSH. For the assay, 50 µL of each diluted sample and standard was dispensed into a 96-well microplate. Subsequently, 25 µL of the Colorimetric Detection Reagent and 25 µL of a reaction mixture containing Glutathione Reductase Solution were added to each well. The plate was then incubated at room temperature for 20 minutes to allow the reaction to occur. Absorbance was measured at 405 nm using a microplate reader. Separate standard curves were constructed for GSH and for 2VP-treated GSSG using serial dilutions of the Glutathione Standard, which enabled the calculation of GSH and GSSG concentrations in the samples. All assays were conducted in duplicate to ensure accuracy, and the results were analyzed to evaluate oxidative stress levels.

### 2.4. Oxidative DNA damage and Antioxidant Enzymes Activity measurement in Plasma by ELISA Assay

The concentration of free 8-hydroxy-2’-deoxyguanosine (8-OHdG) in plasma samples was determined using a competitive ELISA kit provided by Stressmarq Biosciences, Canada (Catalog Number: SKC-120A), following the manufacturer’s protocol as previously described by us [30]. Additionally, the concentrations of Human Cu/ZnSOD in plasma were quantified according to the specific protocols provided by the manufacturers (Invitrogen, USA).

### 2.5. Statistical Analyses

Statistical analyses of oxidative stress and plasma proteomic profiles were conducted using GraphPad Prism version 10.0.3 (San Diego, CA, USA). To ensure baseline similarities and make comparisons between groups, an independent t-test was employed. P-values were determined using one-way ANOVA followed by Tukey’s multiple comparison test while keeping the threshold for statistical significance at p ≤ 0.05. The results are expressed as the mean ± standard deviation (SD).

## Results

### 3.1. Baseline characteristics of the cohort studied

Samples collected during the initial phase of the pandemic (2020-2021) were included in this study and the previous study [28] (**Table 1**). Plasma samples were collected from individuals attending the COVID-19 clinic where SARS-CoV-2 infection was confirmed by RT-PCR as previously described [27]. Un-infected controls (Un-inf) were SARS-CoV-2 RT-PCR negative at the time of sample collection. Convalescent (Conv) and PCC groups were PCR negative by one month after infection. Conv group was free of any infection related symptoms at 3 months post-infection, while the PCC group had persistent symptoms as defined by the WHO criteria prevalent during the sample collection time frame [29].

**Table 1.** Baseline clinical characteristics of study group.

| **Variables** | **No infection**<br>**n = 35** | **Convalescent**<br>**n = 58** | **PCC**<br>**n = 56** |
| --- | --- | --- | --- |
| **Female Sex** |  |  |  |
| n (%) | 21 (60.0) | 23 (39.6) | 37 (66.0) |
| **age range** | 18-73 | 18-81 | 18-88 |
| **Comorbidities (n)** |  |  |  |
| Obesity |  | 9 (15.6) | 9 (15.6) |
| Hypertension | 0 | 1 | 5 |
| Diabetes | 1 | 4 | 7 |
| asthma | 1 | 2 | 6 |
| **number of PCC symptoms**<br>**at 3 mo post-infection**<br>**range (mean)** | 0 | 0 | 1-16 (2.9±2.8) |

PCC- convalescent individuals with PCC; PCC= post-covid condition

### 3.2. Proteomics Analysis

#### 3.2.1. DIA-MS-based differentially expressed proteins and PCA analysis

We used a data-independent acquisition (DIA) proteomics strategy to analyze plasma collected at 3 months post-PCR confirmed infections from 53 individuals with PCC, 62 recovered individuals without PCC and 35 healthy controls (PCR negative). We identified 235 unique proteins from these groups and quantified them using DIA-NN, focusing on prototypic peptides uniquely mapped to specific proteins. PCA analysis (**Fig. 1A**) showed that the convalescent and PCC individuals were distinct from un-infected individuals, but the differences between the convalescent and PCC groups were relatively small, reflecting few differences in protein expression profiles. This was reflected in the Venn diagram (**Fig. 1B**) where 224 unique proteins were commonly found among all the groups, eight proteins were unique to PCC individuals, nine were unique to convalescent individuals and one unique to uninfected individuals. The heatmap analysis distinctly separates the three groups based on unique peptide expression profiles (**Fig. 1C**). The differences observed in the Ig-H, Ig-l and Ig-k peptides may reflect the immunoglobulin usage in the antibody responses to SARS-CoV-2. There is a broad overlap in differentially abundant proteins (DEPs) between PCC and convalescent groups compared to uninfected individuals, indicating that the changes brought about by mild COVID-19 infection persists at least for 3 months and does not does not aid in distinguishing Conv from PCC. (**Fig. 1D**). DEPs analyses showed differential abundance in 49 proteins in the convalescent group compared to un-infected individuals, of which 26 were upregulated and 23 were downregulated (**Fig. 1E (left panel**). Similarly, 53 proteins were differentially expressed between PCC and un-infected while only 2 proteins showed differential expression between PCC and convalescent individuals (**Fig. 1E (middle and right panel**).

**Figure 1.**
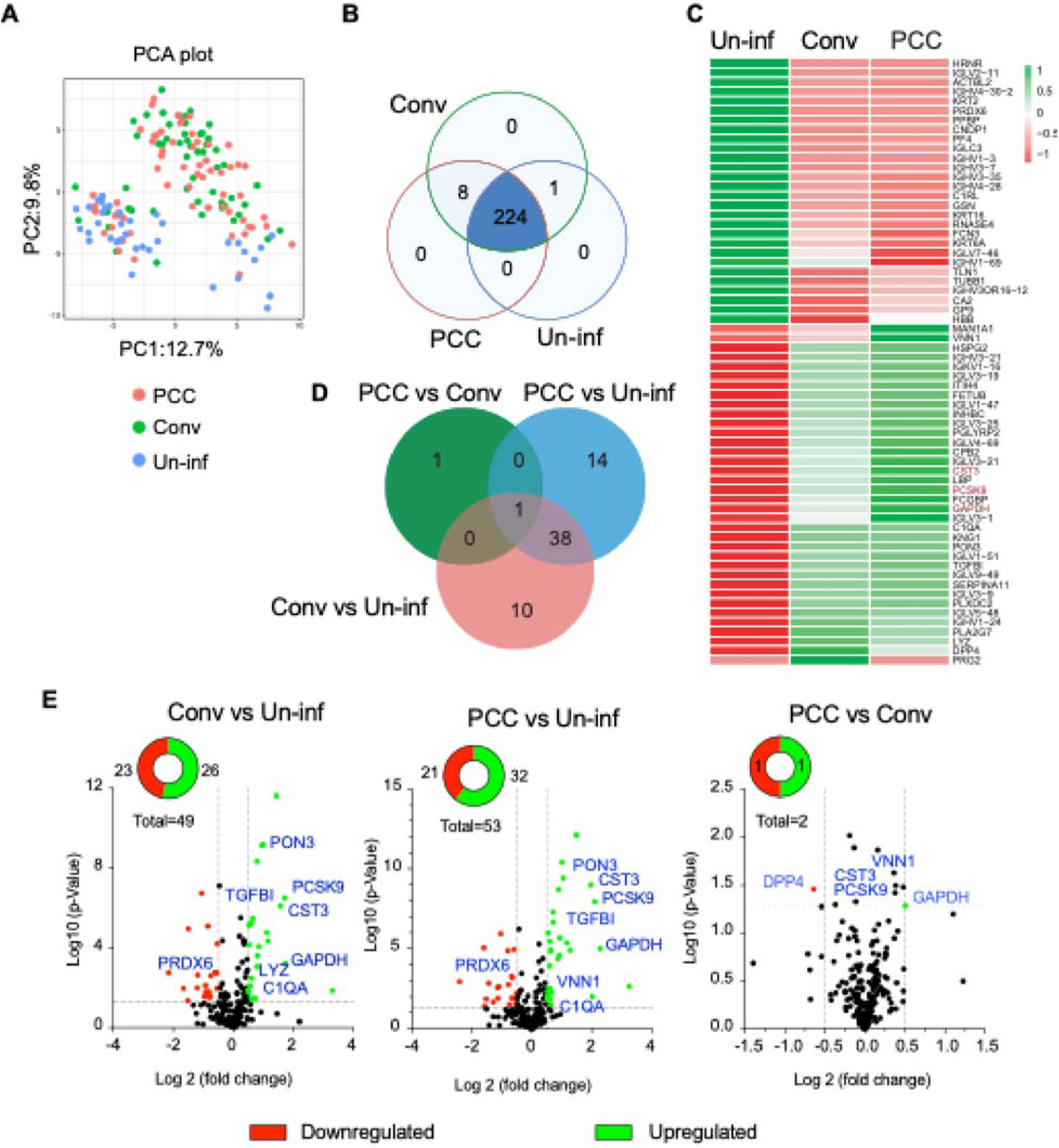
Proteomic analyses. (A) PCA plot analysis. The PCA plot separated the groups into two principal components, PC1 and PC2. Each dot represents an individual sample in the PCA plot. (B) Venn diagram represents each group’s shared common protein and unique proteins, with the overlapping region denoting common proteins across all three conditions. The DIA-Analyst Application platform (Analyst Suites) was used for PCA and Venn diagram analyses. (C) Comprehensive Heatmap analysis for differentially expressed peptide abundance profiles. (D) The Venn diagram shows unique and common significant differential proteins in PCC, convalescent and un-infected individuals, with the central overlap representing the proteins shared among the groups. (E) Comparative Volcano plot analysis of plasma protein expression among three groups. Proteins that are significantly modulated (fold change of +/- 0.5 and p-value of 0.05, indicated by dotted line) are labeled and shown in red and green color. The x-axis represents the log2 fold change, and the y-axis displays the -log10 p-value.

Analyses of protein-protein interaction network by STRING database (version 12.2) among the differentially expressed proteins between PCC and uninfected individuals, showed that 19 proteins formed a single network, whereas eleven differentially expressed proteins were not related (**Fig. 2A**). The resulting data were analyzed using Cytoscape software following a Cyto hub parameter to identify the most influential proteins. The top12 hub proteins are selected based on their degree of connectivity, and those are associated with different functional activities, such as oxidative stress and metabolic activity (GAPDH), inflammation (CST3, TGFB1), and complement and coagulation cascades (C1QA, KNG1, CBP2) (**Fig. 2B**). Similarly, twenty-one (21) differentially expressed protein interactions are observed in a single cluster in the comparison between convalescent individuals and un-infected individuals (**Fig. 2B**). The Cyto hub identified eleven top hubs (**Fig. 2A**) that are similar to those identified in **Fig. 2B**. Overall these findings indicate comparable proteomic profile at 3 months in the SARS-CoV-2 infected group, irrespective of the PCC status.

**Figure 2:**
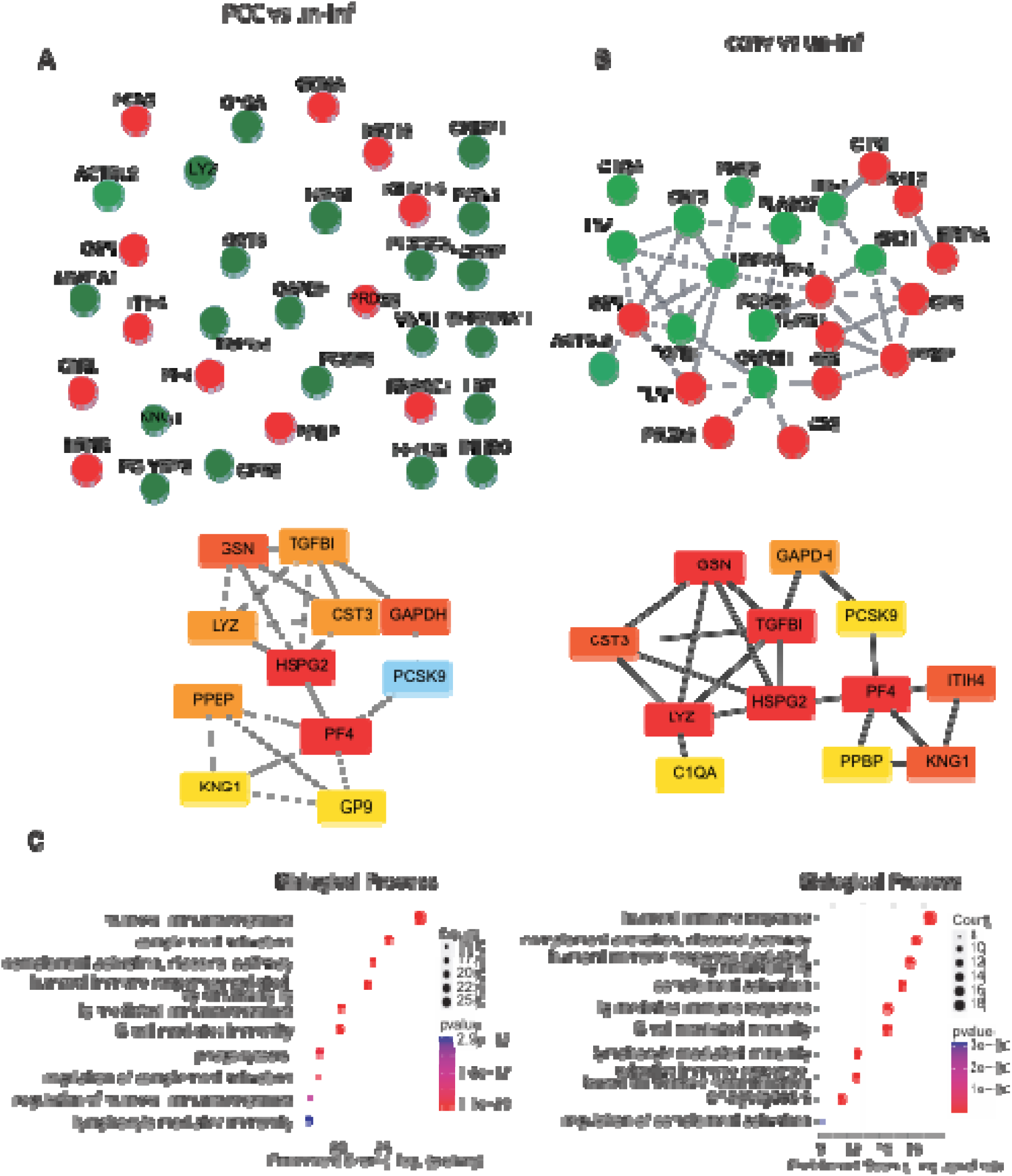
Protein-Protein Interaction Network (PPI) analysis for the differential expressed protein. (A-B) PPI network illustrates significant differentially expressed proteins (21) between PCC (A), convalescent (B) and uninfected individuals. Lower panel highlights the top ten hub proteins identified by CytoHubba in Cytoscape based on the degree of connectivity of each protein node. In this network, nodes represent individual proteins, whereas edges indicate their interactions. The green and red nodes indicate the upregulated and downregulated proteins. The PPI network was constructed using the STRING database version 12.2 and visualized with Cytoscape 3.10.2. C) Comparative Gene Ontology (GO) and KEGG Pathway Analysis.

GO enrichment analyses of the differentially expressed proteins between the PCC, convalescent and uninfected groups showed terms related to immune responses (**Fig. 2C**). Humoral immune response (GO:0006959) and immunoglobulin-mediated immune response (GO:0016064) reflect the differences in the antigen receptor profiles probably resulting from the immune responses generated against SARAS-CoV-2 infection. While the persistence of the differences in the antibody profiles 3 months after infection reflect the long lifespan of circulating antibodies, it was striking that differences in the complement pathways (GO:0006956, GO:0006958) and phagocytosis (GO:0006909) were persisting at 3 months in both the PCC and the convalescent groups. Collectively these observations indicate perturbation in the immune homeostasis well after clearance of the virus.

#### 3.2.2. Validation of differentially expressed proteins or biomarkers by peptide intensity and ELISA assay quantification

To validate the differentially expressed proteins in plasma samples among the three groups, proteins (CST3, GAPDH, PCSK9, C1q, CBPB2, and PRXD6) with significant fold changes and pathway enrichment analysis scores were selected. Given that DIA-NN is an unbiased technique for the quantification of peptide intensities, we sought to validate the presence of possible epitopes of these peptides via ELISA. We randomly selected 40 PCC, 20 Conv and 20 uninfected individuals for validation using commercially available ELISA kits. The GAPDH peptide displays a significant intensity in PCC individuals that is not mimicked in Conv individuals. This trend is recapitulated when GAPDH Ab-epitope detection is assessed via ELISA (**Fig. 3A**). DPP4 was not detected in uninfected samples by proteomics, whereas a fraction of convalescent and PCC samples were positive for DPP4 showing significant differences between the 2 groups (**Fig. 3B**). DPP4 was detected by ELISA in all the 3 groups and the values were comparable. It is a serine protease enzyme that circulates in the blood plasma and is crucial for inflammation and immunity. Three proteins involved in the complement/coagulation pathways showed significant differences in the proteome of PCC. CPB2, a regulator of fibrinolysis, was significantly higher in PCC individuals compared to uninfected individuals by proteomics and in ELISA assays (Fig. 3C). CBPB2, peptide abundance reflected the GAPDH data, but in ELISA both the PCC and convalescent groups were significantly higher than the uninfected group (Fig. 3C). Similarly, peptide abundance for complement component protein C1q was significantly increased PCC and convalescent groups when compared to uninfected individuals, whereas the ELISA results were comparable (Fig. 3D). Although stratification of the data by biological sex indicated some differences, the limited sample size precludes definitive conclusions. Therefore, these findings should be interpreted with caution due to the small number of patients available for this subgroup analysis. (**Supplementary Fig. 1**). Cystatin C (CST3) and Proprotein convertase subtilisin/kexin type 9 (PCSK9) have been shown to be associated with the COVID-19 pathology in various studies [32–34]. CST3, a potent inhibitor of cysteine proteases was highly upregulated in PCC and in convalescent individuals when compared to uninfected controls (Fig. 3E). However, CST3 ELISA demonstrated opposite trend with reduced CST3 levels in PCC and convalescent group relative to uninfected individuals (**Supplementary Fig. 2A**). On the other hand, PCSK9 is significantly increased in PCC and convalescent group when compared to uninfected individuals by peptide abundance but not in ELISA (**Fig. 3F, Supplementary Fig. 2B**). This discrepancy between peptide abundance and ELISA measurements may reflect post-translational modifications, proteolytic processing, or conformational changes in CST3 and PCSK9 that alter epitope accessibility for antibody-based detection, despite increased peptide-level representation observed by mass spectrometry.

**Figure 3:**
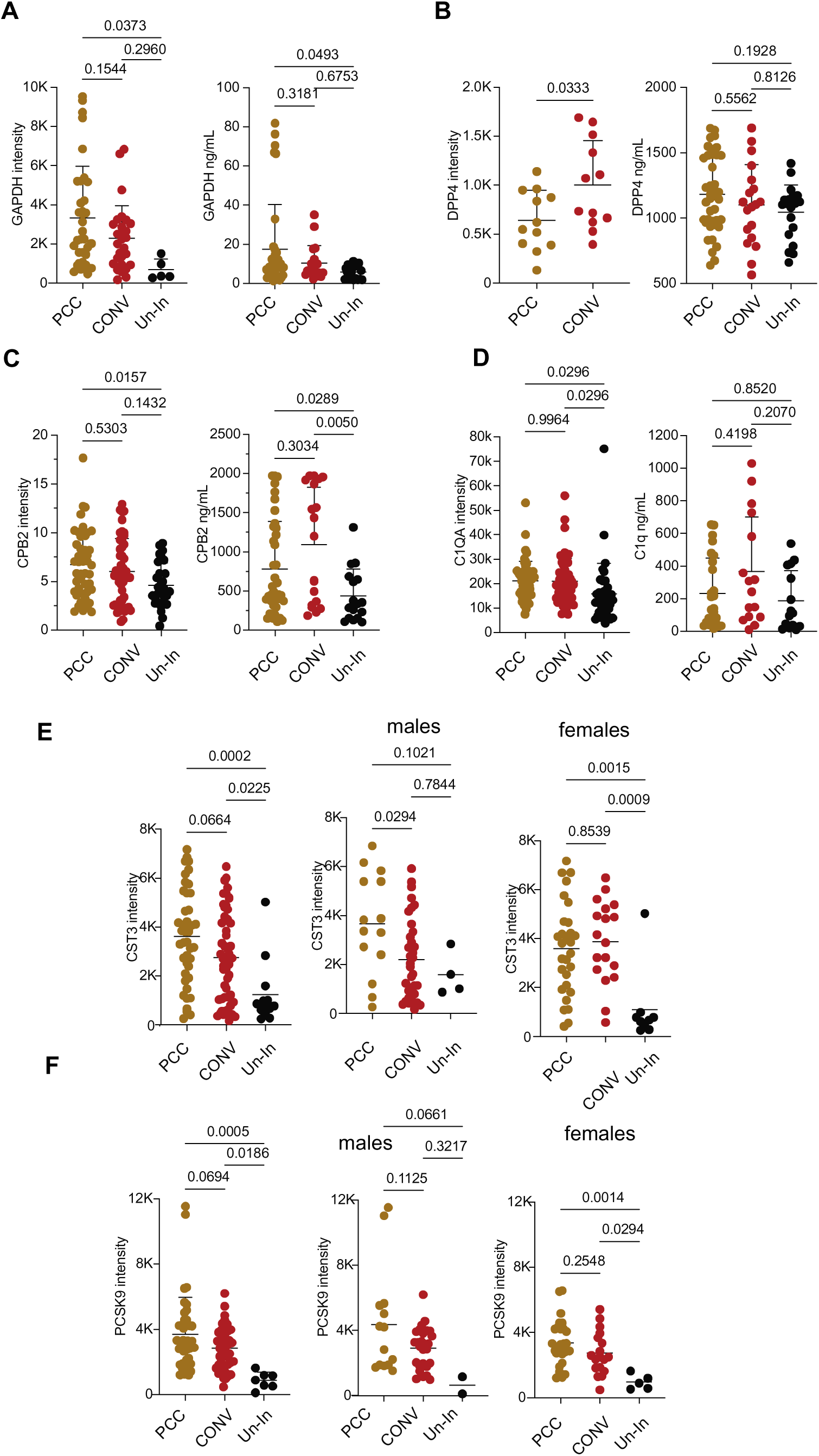
Comparison between proteomics and ELISA measurements for a restricted set of proteins. Peptide abundance (DIA-MS) and protein levels (ELISA) are shown for select proteins across three groups. Statistical analysis was conducted using one-way/two-way ANOVA followed by Tukey’s multiple comparisons tests, with significance set at a p-value ≤ 0.05. Data are presented as mean with standard deviation.

We combined the proteomics and ELISA data to determine whether the PCC group can be separated from the convalescent group using PCA plot (**Fig. 4**). The PCA plot visualizes the combination and differences of peptide intensity and protein detection by ELISA in a 2D plot. The data took into account the six proteins (GAPDH, PCSK9, DPP4, CBPB2, C1q & CST3) that were tested by ELISA. PCC, convalescent and the uninfected groups show overlapping clusters but distinct proteomic profiles, although healthy controls form distinct clusters. The uninfected group clusters tightly in the right quadrant, showing low variability and distinct separation from PCC and convalescent group. The convalescent group is partially overlapping with both PCC and uninfected group suggesting an intermediate molecular profile. The PCC group clusters distinctly to the left, indicating a unique proteomic, distinguishable from the other 2 groups. Arrows show the direction and magnitude of each protein’s influence. Longer arrows suggest stronger contribution to sample separation between the groups. Direction indicates which group(s) the proteins areassociated with. GAPDH, PCSK9, CST3, CPB2, and C1QA are all pointing towards the PCC group, especially their ELISA-detected forms (suffix 2). This suggests higher expression levels or stronger discriminatory power in PCC subjects. ELISA and MS detections are relatively concordant (similar directions), showing consistency across platforms.

**Figure 4:**
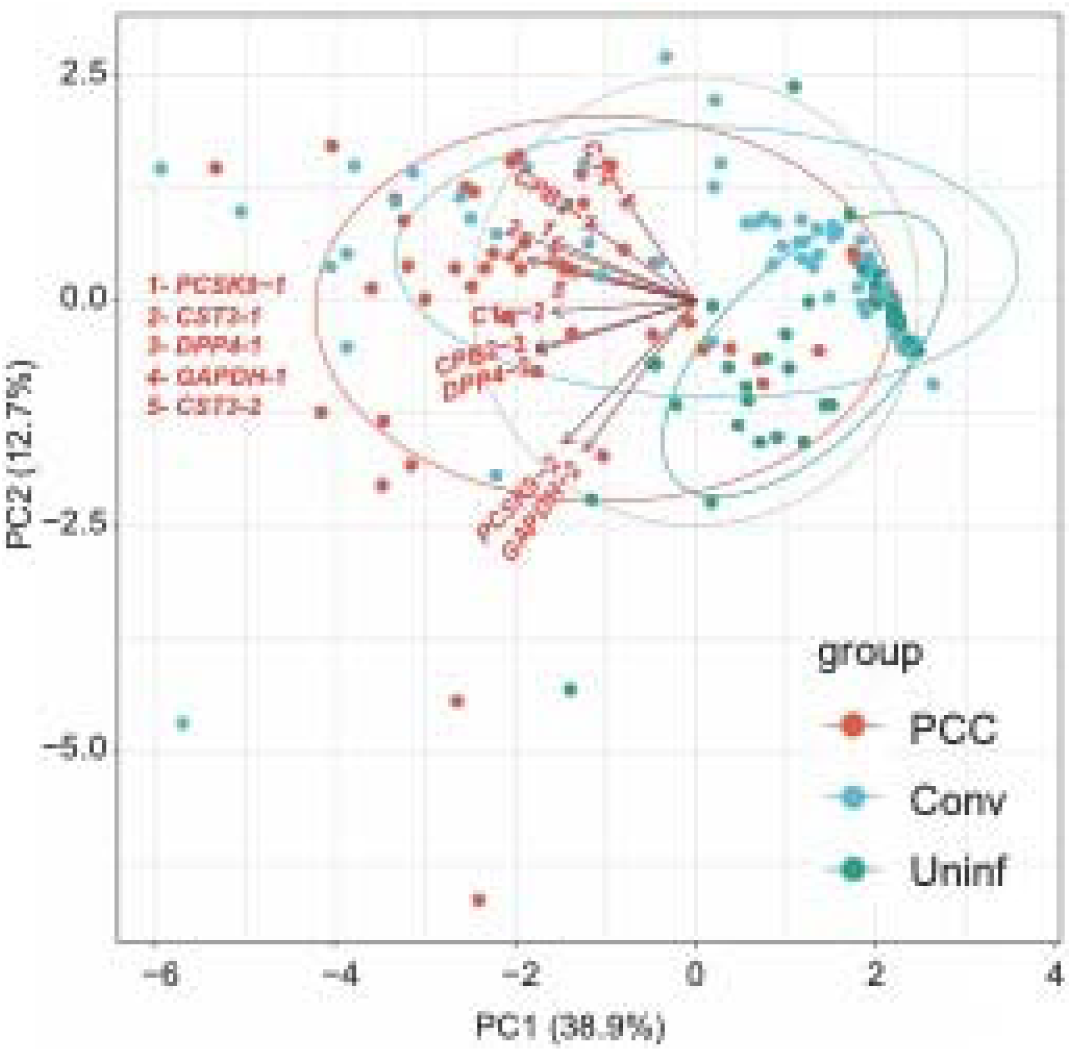
Validation of differential expressed protein by PCA. The PCA plot shows the separation of three groups. The ellipses provide a visual sense of how much variability exists within each group and how much overlap (similarity) or separation (difference) there is between the groups. Less overlap suggests a more distinct separation between the groups, while more overlap suggests similarity. Arrows indicate the weightage of peptides relevant to proteins such as PCSK9, GAPDH, DPP4, CPB2, and CST3 contributing to the separation of the groups along the principal components (PC1 & PC2). In the PCA plot, 1= Peptide intensity by DIA-MS; and 2= intact proteins by ELISA assay

### 3.3. Analysis of Antioxidant activity and DNA-damaged oxidative stress

Various studies have shown the association between SARS-CoV-2 infection and decreased ability to cope with oxidative stress [25, 35]. However, the status of oxidative stress is not well known in the PCC group [26, 36, 37]. In the proteomic analyses we identified few proteins implicated in the oxidative stress pathway that were differentially expressed (**Fig. 5A, Supplementary Fig. 3**). Peroxiredoxin 6 or PRDX6, a bifunctional protein with anti-oxidant property [38] was significantly downregulated in the Conv and PCC groups when compared to uninfected controls. Paraoxonase-3 (PON3) and Vanin1 (VNN1) that are upregulated under conditions of oxidative stress [39, 40], were significantly increased in PCC groups (**Fig. 5B, C**) when compared to uninfected controls. PON3 protects from mitochondrial oxidative stress [41]. VNN1 levels are increased in conditions associated with increased oxidative stress [42]. These observations suggest that compared to uninfected individuals, COVID-19-induced oxidative stress persists 3 months after infection.

**Figure 5:**
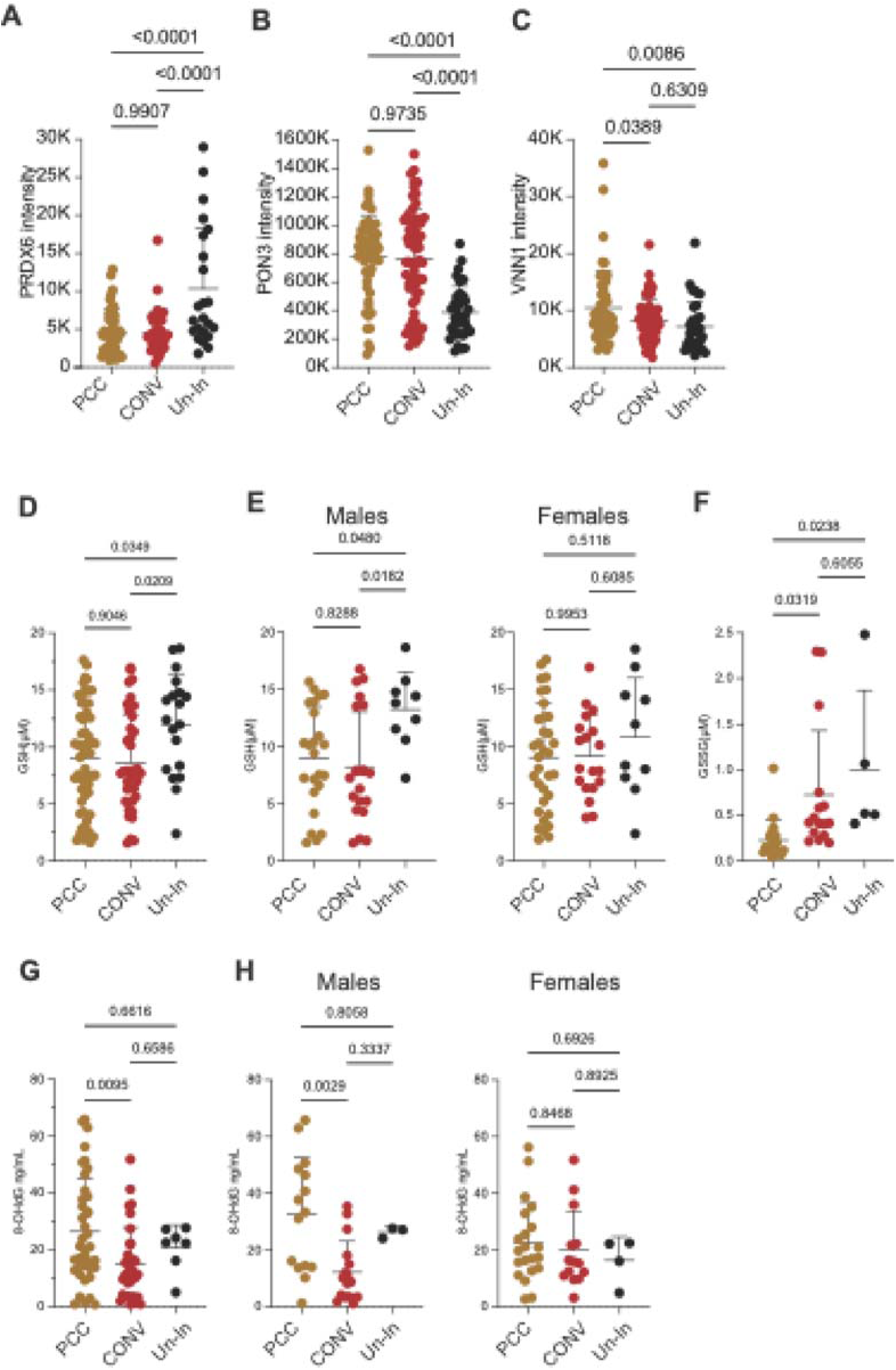
Expression levels of peptides and markers of oxidative stress in plasma. (A-C) Peptide abundance (DIA-MS) for PRDX6, PON3 and VNN; (D-E) GSH, (F) GSSG and (G,H) 8-OHdG DNA concentrations in the three groups. Statistical comparisons were carried out using one-way ANOVA followed by Tukey’s multiple comparisons test with significance determined at a p-value ≤ 0.05. The data are presented as mean + standard deviation (SD), and different colors represent this group. GSH - glutathione; GSSG - Glutathione disulfide; 8-OHdG - 8-hydroxy-2’-deoxyguanosine; PCC – post COVID condition; CONV – convalescent; Un-In – uninfected.

In addition to the proteomic approach, we measured the GSH levels in the plasma samples at 3 months post infection in PCC and Conv groups. No significant difference in GSH levels was observed between the between the PCC and convalescent groups (**Fig. 5D**). In contrast, both groups exhibited significantly lower GSH concentrations compared to the uninfected controls (p ≥ 0.05). When the analysis is segregated into biological male and female groups, PCC and convalescent males showed significantly reduced levels of GSH (**Fig. 5E**) compared to uninfected males whereas no noticeable differences are observed in female groups. These observations suggest a possible depletion of GSH during and following mild COVID-19 infection that appear to persist irrespective of PCC. On the other hand, the levels of oxidized GSSG (**Fig. 5F**) were significantly lower in the PCC compared to the convalescent group (p ≥ 0.05). These data suggest that oxidized GSH levels and total GSH levels show differential recovery at 3 months following COVID-19 infection and may have different profiles between PCC and convalescent groups. We assessed the concentration of other anti-oxidant enzyme Cu/Zn SOD, in the plasma by ELISA but its level was comparable between the 3 groups (**Supplementary Fig. 4**). 8-OHdG is produced when reactive oxygen species (ROS), specifically hydroxyl radicals, oxidize the guanine base in DNA, indicating oxidative DNA damage, that can be detected in circulation. 8-OHdG levels were significantly elevated in the PCC group when compared to the convalescent group (**Fig. 5G**). Segregation by biological sex showed significant increases in males but not in females with PCC when compared to the convalescent group, reflecting the results of GSH (**Fig. 5H**).

## Discussion

Our study shows that the proteome of plasma from SARS-CoV-2 naïve group is distinct from that of convalescent and PCC group. These results were surprising as we expected convalescent individuals to be similar to that of uninfected individuals. With the goal of identifying plasma biomarkers by proteomics, various studies have analyzed cohorts involving patients with severe COVID-19 disease [43–49], while studies on patients with mild infections that comprise a majority of the long COVID patients are few [50]. The plasma proteome of uninfected individuals is distinct from that of convalescent and PCC groups even after mild infection. It is not clear if this observation is unique to COVID-19 or can be generalized to other viral infections. Furthermore, 3 months following mild COVID-19 infection, the differences in the plasma proteome are subtle between the two infected groups. In this paper the relevance of mass-spec-based techniques for biomarker analyses has been strengthened through partial validation by ELISA. Our results confirm the accuracy of the proteomic data in both quantification methods using a small cohort. However, only two proteins out of six, namely GAPDH and CPB2, are more consistent in both analyses, demonstrating that the biomarkers identified in this study hold significant promise for clinical applications in monitoring post-COVID condition (PCC) progression, while the rest vary too much between assays to be reliable.

The involvement of oxidative stress in PCC has been addressed by various groups [22, 23, 26, 51–55]. Lower levels of circulating free thiols persisted as a marker of oxidative stress in non-hospitalized infected individuals who when on to develop long COVID [23]. In line with these observations, our omics and oxidative stress quantification data indicate that at 3 months post infection, convalescent individuals with or without PCC exhibit alterations in the markers of oxidative stress that are significantly different from uninfected individuals. Comparable analyses over time will validate the relevance of these markers in PCC patients.

Long COVID or PCC encompasses a wide spectrum of symptoms with varying degree of intensity that has limited predictable correlation with the SARS-CoV-2 infection severity [56]. Five years after the global spread of COVID-19, persistence of PCC has debilitating effects on long COVID patients and compromises their quality of life. Longitudinal studies suggest that an increase in number of symptoms during infection as a plausible predictor of PCC [57–59]. We and others have shown that PCC is characterized by uncoordinated immune responses to SARS-CoV-2 antigens [28, 60]. In line with these observations other groups have identified altered immune profiles in PCC [50]. Inflammation related biomarkers and autoantibodies have been observed to be increased in PCC patients when compared to convalescent individuals following severe infections [47, 61]. Different approaches, different techniques strongly suggest that long COVID can be distinguished by specific marker, but would require the integration of the clinical spectrum of long COVID symptoms and the degree of the phenotype as a disease.

### Limitations and strengths of the study and future research

The obvious limitation of this study is the small sample size. We restricted ourselves to non-hospitalized individuals during the early phase of the pandemic between 2020 and 2021, when infection in the general population was well controlled due to severe restrictions. While the diagnosis of PCC was based on the 2020 criteria [29], PCC incidence and associated predominant symptoms may be dependent on the variants that were in circulation [62]. Our study also provides an important outcome that oxidative stress can be monitored by minimally invasive methods of analyses of the associated biological processes in the peripheral blood. Measurement of different parameters related to oxidative stress would have provided a complete picture.

## Author Contributions

Study design and funding: SR, AP, HAC and SI; identification of clinical samples: CR-P and AP; sample preparation for proteomics and data analyses: MMHC, JFL; Oxidative stress: MMHC and SI; ELISA: MMHC and AJIQ; data compilation and analyses: MMHC and SR; manuscript writing: MMHC, AP, and SR; Manuscript editing, reviewing: MMHC, AJIQ, SR, AP and HAC.

## Funding

This work was supported by the Canadian Institutes of Health Research pandemic response (GA4-177773 and CIHR Pandemic Preparedness and Health Emergencies Research funding (202309PPE-512214) to SR and AP). H.A.C. is a Junior 1 Clinical Research Scholar from the Fonds de recherche du Québec-Santé and holds the André-Lussier research chair of the Université de Sherbrooke. MMHC is the recipient of PhD scholarship from the Faculty of Medicine, Université de Sherbrooke.

## Institutional Review Board Statement

The human studies were approved by the Institutional human ethics committee, CIUSSS de l’Estrie – CHUS, Université de Sherbrooke (approval number 2022-4415).

## Informed Consent Statement

Informed consent was obtained from all subjects involved in the study.

## Data Availability Statement

The data can be made available after obtaining relevant authorizations.

## Conflicts of Interest

HAC received research funds from Janssen, Eli Lilly, Fresenius Kabi and Pfizer, but nothing related to the current manuscript. He also participated in medical education of Pfizer which was conducing trial on PCC, but nothing directly related to the current manuscript or that could impact our interpretation of data. The other authors declare no conflict of interest.

## Notes

### Competing Interest Statement

The authors have declared no competing interest.

## References

1. Pavone P, Ceccarelli M, Taibi R, La Rocca G, Nunnari G: Outbreak of COVID-19 infection in children: fear and serenity. Eur Rev Med Pharmacol Sci 2020, 24(8):4572–4575.

2. Korean Society of Infectious D, Korean Society of Pediatric Infectious D, Korean Society of E, Korean Society for Antimicrobial T, Korean Society for Healthcare-associated Infection C, Prevention, Korea Centers for Disease C, Prevention: Report on the Epidemiological Features of Coronavirus Disease 2019 (COVID-19) Outbreak in the Republic of Korea from January 19 to March 2, 2020. J Korean Med Sci 2020, 35(10):e112.

3. Mahase E: Covid-19: What do we know about “long covid”? BMJ 2020, 370:m2815.

4. Carfi A, Bernabei R, Landi F, Gemelli Against C-P-ACSG: Persistent Symptoms in Patients After Acute COVID-19. JAMA 2020, 324(6):603–605.

5. Dennis A, Wamil M, Alberts J, Oben J, Cuthbertson DJ, Wootton D, Crooks M, Gabbay M, Brady M, Hishmeh L et al: Multiorgan impairment in low-risk individuals with post-COVID-19 syndrome: a prospective, community-based study. BMJ Open 2021, 11(3):e048391.

6. Xie Y, Bowe B, Al-Aly Z: Burdens of post-acute sequelae of COVID-19 by severity of acute infection, demographics and health status. Nat Commun 2021, 12(1):6571.

7. Davis HE, McCorkell L, Vogel JM, Topol EJ: Long COVID: major findings, mechanisms and recommendations. Nat Rev Microbiol 2023, 21(3):133–146.

8. Ballering AV, van Zon SKR, Olde Hartman TC, Rosmalen JGM, Lifelines Corona Research I: Persistence of somatic symptoms after COVID-19 in the Netherlands: an observational cohort study. Lancet 2022, 400(10350):452–461.

9. Meeting the challenge of long COVID. Nat Med 2020, 26(12):1803.

10. Buonsenso D, Munblit D, De Rose C, Sinatti D, Ricchiuto A, Carfi A, Valentini P: Preliminary Evidence on Long Covid in children. Acta Paediatr 2021.

11. Global Burden of Disease Long CC, Wulf Hanson S, Abbafati C, Aerts JG, Al-Aly Z, Ashbaugh C, Ballouz T, Blyuss O, Bobkova P, Bonsel G et al: Estimated Global Proportions of Individuals With Persistent Fatigue, Cognitive, and Respiratory Symptom Clusters Following Symptomatic COVID-19 in 2020 and 2021. JAMA 2022, 328(16):1604–1615.

12. Geng LN, Erlandson KM, Hornig M, Letts R, Selvaggi C, Ashktorab H, Atieh O, Bartram L, Brim H, Brosnahan SB et al: 2024 Update of the RECOVER-Adult Long COVID Research Index. JAMA 2025, 333(8):694–700.

13. Mizrahi B, Sudry T, Flaks-Manov N, Yehezkelli Y, Kalkstein N, Akiva P, Ekka-Zohar A, Ben David SS, Lerner U, Bivas-Benita M et al: Long covid outcomes at one year after mild SARS-CoV-2 infection: nationwide cohort study. BMJ 2023, 380:e072529.

14. Nalbandian A, Sehgal K, Gupta A, Madhavan MV, McGroder C, Stevens JS, Cook JR, Nordvig AS, Shalev D, Sehrawat TS et al: Post-acute COVID-19 syndrome. Nat Med 2021.

15. Huang C, Wang Y, Li X, Ren L, Zhao J, Hu Y, Zhang L, Fan G, Xu J, Gu X et al: Clinical features of patients infected with 2019 novel coronavirus in Wuhan, China. Lancet 2020, 395(10223):497–506.

16. Bellan M, Soddu D, Balbo PE, Baricich A, Zeppegno P, Avanzi GC, Baldon G, Bartolomei G, Battaglia M, Battistini S et al: Respiratory and Psychophysical Sequelae Among Patients With COVID-19 Four Months After Hospital Discharge. JAMA Netw Open 2021, 4(1):e2036142.

17. Sykes DL, Holdsworth L, Jawad N, Gunasekera P, Morice AH, Crooks MG: Post-COVID-19 Symptom Burden: What is Long-COVID and How Should We Manage It? Lung 2021, 199:113–119.

18. Sudre CH, Murray B, Varsavsky T, Graham MS, Penfold RS, Bowyer RC, Pujol JC, Klaser K, Antonelli M, Canas LS et al: Attributes and predictors of long COVID. Nat Med 2021, 27:626–631.

19. Kamal M, Abo Omirah M, Hussein A, Saeed H: Assessment and characterisation of post-COVID-19 manifestations. Int J Clin Pract 2020:e13746.

20. Soriano JB, Murthy S, Marshall JC, Relan P, Diaz JV, Condition WHOCCDWGoP-C-: A clinical case definition of post-COVID-19 condition by a Delphi consensus. Lancet Infect Dis 2022, 22(4):e102–e107.

21. Mehandru S, Merad M: Pathological sequelae of long-haul COVID. Nat Immunol 2022, 23(2):194–202.

22. Stufano A, Isgro C, Palese LL, Caretta P, De Maria L, Lovreglio P, Sardanelli AM: Oxidative Damage and Post-COVID Syndrome: A Cross-Sectional Study in a Cohort of Italian Workers. Int J Mol Sci 2023, 24(8).

23. Vlaming-van Eijk LE, Bulthuis MLC, van der Gun BTF, Wold KI, Veloo ACM, Vincenti Gonzalez MF, de Borst MH, den Dunnen WFA, Hillebrands JL, van Goor H et al: Systemic oxidative stress associates with the development of post-COVID-19 syndrome in non-hospitalized individuals. Redox Biol 2024, 76:103310.

24. Pincemail J, Cavalier E, Charlier C, Cheramy-Bien JP, Brevers E, Courtois A, Fadeur M, Meziane S, Goff CL, Misset B et al: Oxidative Stress Status in COVID-19 Patients Hospitalized in Intensive Care Unit for Severe Pneumonia. A Pilot Study. Antioxidants (Basel) 2021, 10(2).

25. Neves FF, Pott-Junior H, Yamashita KMC, de Sousa Santos S, Cominetti MR, de Melo Freire CC, Cunha AFD, Jordao Junior AA: Do the oxidative stress biomarkers predict COVID-19 outcome? An in-hospital cohort study. Free Radic Biol Med 2023, 207:194–199.

26. Valente Coronel PM, Luiz Soares Basilio DC, Teixeira Espinoca I, Souza de Souza KF, Miranda Campos N, Seiji Nakano Ota R, Paredes-Gamero EJ, Wilhelm Filho D, Coimbra Motta-Castro AR, Trentin Perdomo R et al: Involvement of oxidative stress in post-acute sequelae of COVID-19: clinical implications. Redox Rep 2025, 30(1):2471738.

27. Tremblay K, Rousseau S, Zawati MH, Auld D, Chasse M, Coderre D, Falcone EL, Gauthier N, Grandvaux N, Gros-Louis F et al: The Biobanque quebecoise de la COVID-19 (BQC19)-A cohort to prospectively study the clinical and biological determinants of COVID-19 clinical trajectories. PLoS One 2021, 16(5):e0245031.

28. Limoges MA, Quenum AJI, Chowdhury MMH, Rexhepi F, Namvarpour M, Akbari SA, Rioux-Perreault C, Nandi M, Lucier JF, Lemaire-Paquette S et al: SARS-CoV-2 spike antigen-specific B cell and antibody responses in pre-vaccination period COVID-19 convalescent males and females with or without post-covid condition. Front Immunol 2023, 14:1223936.

29. WHO: A clinical case definition of post COVID-19 condition by a Delphi consensus. October 6, 2021. https://www.who.int/publications-detail-redirect/WHO-2019-nCoV-Post_COVID-19_condition-Clinical_case_definition-20211 2021.

30. Chowdhury MMH, Fontaine MN, Lord SE, Quenum AJI, Limoges MA, Rioux-Perreault C, Lucier JF, Cliche DO, Levesque D, Boisvert FM et al: Impact of a tailored exercise regimen on physical capacity and plasma proteome profile in post-COVID-19 condition. Front Physiol 2024, 15:1416639.

31. Demichev V, Messner CB, Vernardis SI, Lilley KS, Ralser M: DIA-NN: neural networks and interference correction enable deep proteome coverage in high throughput. Nat Methods 2020, 17(1):41–44.

32. Chattopadhyay P, Khare K, Kumar M, Mishra P, Anand A, Maurya R, Gupta R, Sahni S, Gupta A, Wadhwa S et al: Single-cell multiomics revealed the dynamics of antigen presentation, immune response and T cell activation in the COVID-19 positive and recovered individuals. Front Immunol 2022, 13:1034159.

33. Chen D, Sun W, Li J, Wei B, Liu W, Wang X, Song F, Chen L, Yang J, Yu L: Serum Cystatin C and Coronavirus Disease 2019: A Potential Inflammatory Biomarker in Predicting Critical Illness and Mortality for Adult Patients. Mediators Inflamm 2020, 2020:3764515.

34. Mester P, Amend P, Schmid S, Muller M, Buechler C, Pavel V: Plasma Proprotein Convertase Subtilisin/Kexin Type 9 (PCSK9) as a Possible Biomarker for Severe COVID-19. Viruses 2023, 15(7).

35. Liu X, Chen R, Li B, Zhang J, Liu P, Li B, Li F, Zhang W, Lyu X, Hu M: Oxidative stress indexes as biomarkers of the severity in COVID-19 patients. Int J Med Sci 2024, 21(15):3034–3045.

36. Kankaya S, Yavuz F, Tari A, Aygun AB, Gunes EG, Bektan Kanat B, Ulugerger Avci G, Yavuzer H, Dincer Y: Glutathione-related antioxidant defence, DNA damage, and DNA repair in patients suffering from post-COVID conditions. Mutagenesis 2023, 38(4):216–226.

37. Erlandson KM, Geng LN, Selvaggi CA, Thaweethai T, Chen P, Erdmann NB, Goldman JD, Henrich TJ, Hornig M, Karlson EW et al: Differentiation of Prior SARS-CoV-2 Infection and Postacute Sequelae by Standard Clinical Laboratory Measurements in the RECOVER Cohort. Ann Intern Med 2024, 177(9):1209–1221.

38. Fisher AB: Peroxiredoxin 6: a bifunctional enzyme with glutathione peroxidase and phospholipase A(2) activities. Antioxid Redox Signal 2011, 15(3):831–844.

39. Mohammed CJ, Lamichhane S, Connolly JA, Soehnlen SM, Khalaf FK, Malhotra D, Haller ST, Isailovic D, Kennedy DJ: A PON for All Seasons: Comparing Paraoxonase Enzyme Substrates, Activity and Action including the Role of PON3 in Health and Disease. Antioxidants (Basel) 2022, 11(3).

40. Zhang B, Lo C, Shen L, Sood R, Jones C, Cusmano-Ozog K, Park-Snyder S, Wong W, Jeng M, Cowan T et al: The role of vanin-1 and oxidative stress-related pathways in distinguishing acute and chronic pediatric ITP. Blood 2011, 117(17):4569–4579.

41. Priyanka K, Singh S, Gill K: Paraoxonase 3: Structure and Its Role in Pathophysiology of Coronary Artery Disease. Biomolecules 2019, 9(12).

42. Yu H, Cui Y, Guo F, Zhu Y, Zhang X, Shang D, Dong D, Xiang H: Vanin1 (VNN1) in chronic diseases: Future directions for targeted therapy. Eur J Pharmacol 2024, 962:176220.

43. Gu X, Wong CCL, Cao B: Authors’ reply to Letter regarding “Probing Long COVID through a Proteomic Lens: a Comprehensive Two-Year Longitudinal Cohort Study of Hospitalised Survivor”. EBioMedicine 2024, 101:105029.

44. Gu X, Wang S, Zhang W, Li C, Guo L, Wang Z, Li H, Zhang H, Zhou Y, Liang W et al: Probing long COVID through a proteomic lens: a comprehensive two-year longitudinal cohort study of hospitalised survivors. EBioMedicine 2023, 98:104851.

45. Yang C, Shannon CP, Tebbutt SJ: Unravelling long COVID: insights from proteomics and considerations for comprehensive understanding. EBioMedicine 2024, 101:105023.

46. Wang K, Khoramjoo M, Srinivasan K, Gordon PMK, Mandal R, Jackson D, Sligl W, Grant MB, Penninger JM, Borchers CH et al: Sequential multi-omics analysis identifies clinical phenotypes and predictive biomarkers for long COVID. Cell Rep Med 2023, 4(11):101254.

47. Canderan G, Muehling LM, Kadl A, Ladd S, Bonham C, Cross CE, Lima SM, Yin X, Sturek JM, Wilson JM et al: Distinct type 1 immune networks underlie the severity of restrictive lung disease after COVID-19. Nat Immunol 2025, 26(4):595–606.

48. Cervia-Hasler C, Bruningk SC, Hoch T, Fan B, Muzio G, Thompson RC, Ceglarek L, Meledin R, Westermann P, Emmenegger M et al: Persistent complement dysregulation with signs of thromboinflammation in active Long Covid. Science 2024, 383(6680):eadg7942.

49. Wolday D, Gebrehiwot AG, Le Minh AN, Rameto MA, Abdella S, Gebreegziabxier A, Amogne W, Rinke de Wit TF, Hailu M, Tollera G et al: Distinct proteomic signatures in Ethiopians predict acute and long-term sequelae of COVID-19. Front Immunol 2025, 16:1575135.

50. Gao Y, Cai C, Adamo S, Biteus E, Kamal H, Dager L, Miners KL, Llewellyn-Lacey S, Ladell K, Amratia PS et al: Identification of soluble biomarkers that associate with distinct manifestations of long COVID. Nat Immunol 2025, 26(5):692–705.

51. Shankar V, Wilhelmy J, Curtis EJ, Michael B, Cervantes L, Mallajosyula VA, Davis RW, Snyder M, Younis S, Robinson WH et al: Oxidative Stress is a shared characteristic of ME/CFS and Long COVID. bioRxiv 2024.

52. Mrakic-Sposta S, Vezzoli A, Garetto G, Paganini M, Camporesi E, Giacon TA, Dellanoce C, Agrimi J, Bosco G: Hyperbaric Oxygen Therapy Counters Oxidative Stress/Inflammation-Driven Symptoms in Long COVID-19 Patients: Preliminary Outcomes. Metabolites 2023, 13(10).

53. Mikuteit M, Baskal S, Klawitter S, Dopfer-Jablonka A, Behrens GMN, Muller F, Schroder D, Klawonn F, Steffens S, Tsikas D: Amino acids, post-translational modifications, nitric oxide, and oxidative stress in serum and urine of long COVID and ex COVID human subjects. Amino Acids 2023, 55(9):1173–1188.

54. Al-Hakeim HK, Al-Rubaye HT, Al-Hadrawi DS, Almulla AF, Maes M: Long-COVID post-viral chronic fatigue and affective symptoms are associated with oxidative damage, lowered antioxidant defenses and inflammation: a proof of concept and mechanism study. Mol Psychiatry 2023, 28(2):564–578.

55. Saleh MG, Chang L, Liang H, Ryan MC, Cunningham E, Garner J, Wilson E, Levine AR, Kottilil S, Ernst T: Ongoing oxidative stress in individuals with post-acute sequelae of COVID-19. NeuroImmune Pharm Ther 2023, 2(2):89–94.

56. Robineau O, Hue S, Surenaud M, Lemogne C, Dorival C, Wiernik E, Brami S, Nicol J, de Lamballerie X, Blanche H et al: Symptoms and pathophysiology of post-acute sequelae following COVID-19 (PASC): a cohort study. EBioMedicine 2025, 117:105792.

57. Subramanian A, Nirantharakumar K, Hughes S, Myles P, Williams T, Gokhale KM, Taverner T, Chandan JS, Brown K, Simms-Williams N et al: Symptoms and risk factors for long COVID in non-hospitalized adults. Nat Med 2022, 28(8):1706–1714.

58. Tanguay P, Decary S, Lemaire-Paquette S, Leonard G, Piche A, Dubois MF, Kairy D, Bravo G, Corriveau H, Marquis N et al: Trajectories of health-related quality of life and their predictors in adult COVID-19 survivors: A longitudinal analysis of the Biobanque Quebecoise de la COVID-19 (BQC-19). Qual Life Res 2023, 32(9):2707–2717.

59. Huang L, Li X, Gu X, Zhang H, Ren L, Guo L, Liu M, Wang Y, Cui D, Wang Y et al: Health outcomes in people 2 years after surviving hospitalisation with COVID-19: a longitudinal cohort study. Lancet Respir Med 2022, 10(9):863–876.

60. Yin K, Peluso MJ, Luo X, Thomas R, Shin MG, Neidleman J, Andrew A, Young KC, Ma T, Hoh R et al: Long COVID manifests with T cell dysregulation, inflammation and an uncoordinated adaptive immune response to SARS-CoV-2. Nat Immunol 2024, 25(2):218–225.

61. Liew F, Efstathiou C, Fontanella S, Richardson M, Saunders R, Swieboda D, Sidhu JK, Ascough S, Moore SC, Mohamed N et al: Large-scale phenotyping of patients with long COVID post-hospitalization reveals mechanistic subtypes of disease. Nat Immunol 2024, 25(4):607–621.

62. Takaoka H, Kawada I, Hiruma G, Nagashima K, Terai H, Ishida N, Namkoong H, Asakura T, Masaki K, Miyata J et al: Long COVID among the first three waves of COVID-19 in Japan: a multicentre cohort study. BMJ Open Respir Res 2024, 11(1).

