## Supplemental Figure 1 for "Molecular Insights into Long COVID: Plasma Proteomics Reveals Oxidative Stress, Coagulation Cascade Activation, and Glycolytic Imbalance"

**Supplementary figures:**

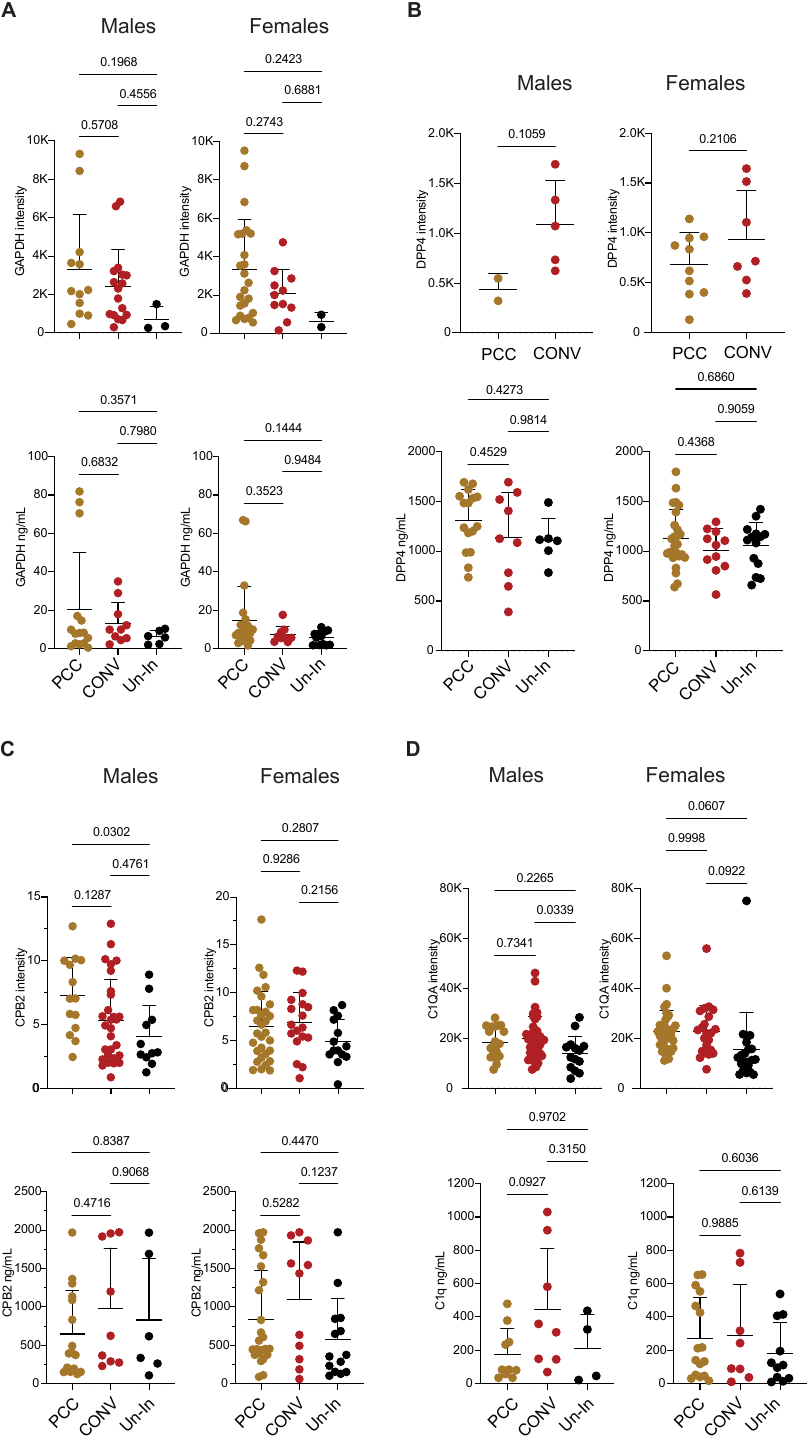

**Supplementary Fig. 1** : **Comparison between proteomics and ELISA measurements for a restricted set of proteins in males and females (at birth).** Peptide abundance (DIA-MS) and protein levels (ELISA) are shown for select proteins across three groups. Statistical analysis was conducted using one-way/two-way ANOVA followed by Tukey's multiple comparisons tests, with significance set at a p-value $\leq$ 0.05. Data are presented as mean with standard deviation.

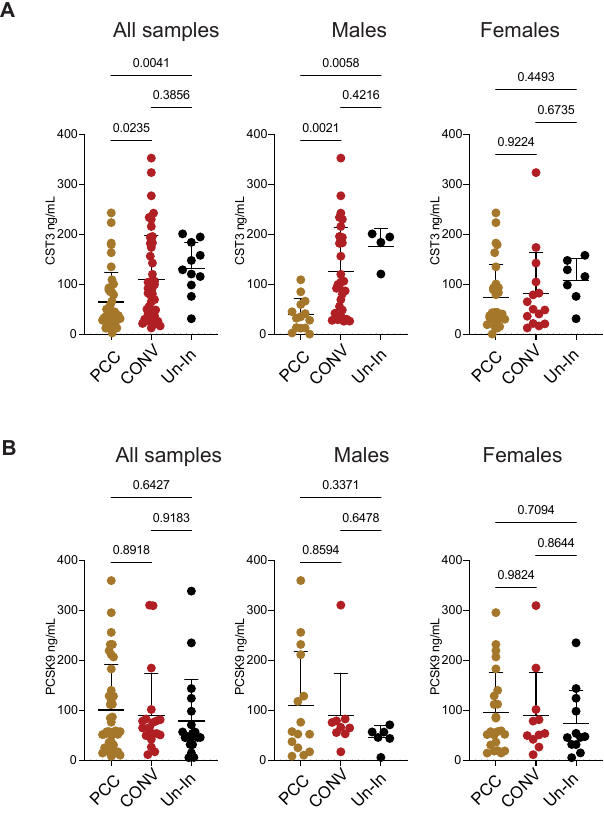

**Supplementary Fig. 2**: **ELISA measurements for a restricted set of proteins in all samples and following segregation into males and females (at birth).** Protein levels (ELISA) are shown for select proteins across three groups. Statistical analysis was conducted using one-way/two-way ANOVA followed by Tukey's multiple comparisons tests, with significance set at a p-value $\leq$ 0.05. Data are presented as mean with standard deviation.

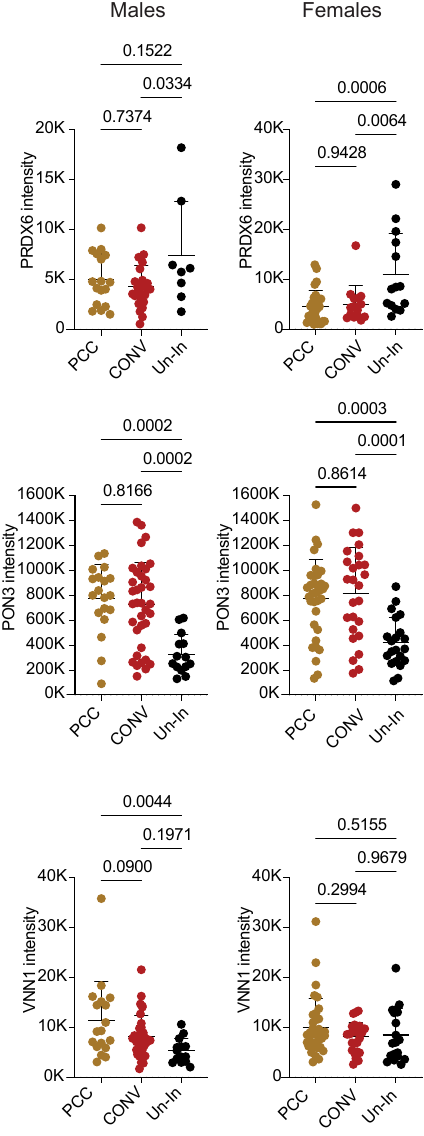

**Supplementary Fig. 3**: **Peptide abundance for oxidative stress related proteins in males and females (at birth).** Peptide abundance (DIA-MS) is shown for select proteins. Statistical analysis was conducted using one-way/two-way ANOVA followed by Tukey's multiple comparisons tests, with significance set at a p-value $\leq$ 0.05. Data are presented as mean with standard deviation.

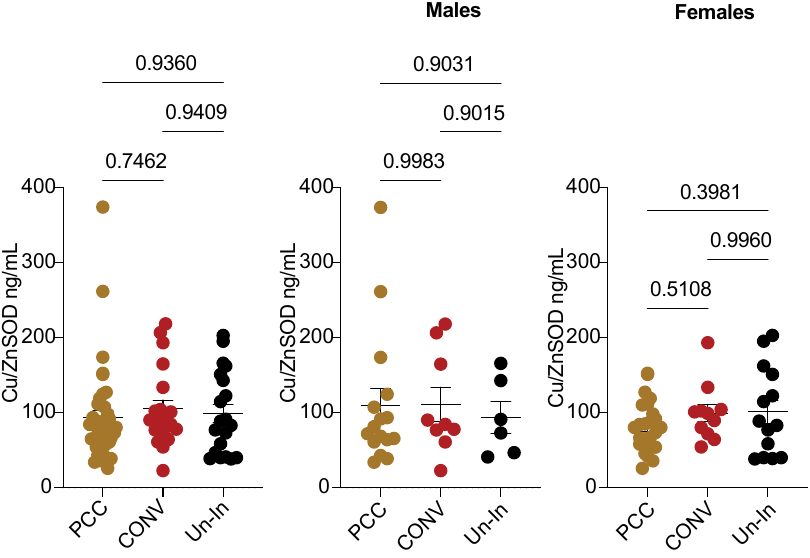

**Supplementary Fig. 4**: **ELISA measurements for Cu/ZnSOD.** Protein levels (ELISA) are shown in all samples and in males and females (at birth). Statistical analysis was conducted using one-way/two-way ANOVA followed by Tukey's multiple comparisons tests, with significance set at a p-value $\leq$ 0.05. Data are presented as mean with standard deviation.
